# Boundary-Forest Clustering: Large-Scale Consensus Clustering of Biological Sequences

**DOI:** 10.1101/2020.04.28.065870

**Authors:** Defne Surujonu, José Bento, Tim van Opijnen

## Abstract

Bacterial species with large sequence diversity enable studies focused on comparative genomics, population genetics and pan-genome evolution. In such analyses it is key to determine whether sequences (e.g. genes) from different strains, are the same or different. This is often achieved by clustering orthologous genes based on sequence similarity. Importantly, one limitation of existing pan-genome clustering methods is that they do not assign a confidence score to the identified clusters. Given that clustering ground truth is unavailable when working with pan-genomes, the absence of confidence scores makes performance evaluation on real data an open challenge. Moreover, most pan-genome clustering solutions do not accommodate cluster augmentation, which is the addition of new sequences to an already clustered set of sequences. Finally, the pan-genome size of many organisms prevents direct application of powerful clustering techniques that do not scale to large datasets. Here, we present Boundary-Forest Clustering (BFClust), a method that addresses these challenges in three main steps: 1) The approximate-nearest-neighbor retrieval method Boundary-Forest is used as a representative selection step; 2) Downstream clustering of the representatives is performed using Markov Clustering (MCL); 3) Consensus clustering is applied across the Boundary-Forest, improving clustering accuracy and enabling confidence score calculation. First, MCL is favorably benchmarked against 6 powerful clustering methods. To explore the strengths of the entire BFClust approach, it is applied to 4 different datasets of the bacterial pathogen *Streptococcus pneumoniae*, and compared against 4 other pan-genome clustering tools. Unlike existing approaches, BFClust is fast, accurate, robust to noise and allows augmentation. Moreover, BFClust uniquely identifies low-confidence clusters in each dataset, which can negatively impact downstream analyses and interpretation of pan-genomes. Being the first tool that outputs confidence scores both when clustering *de novo*, and during cluster augmentation, BFClust offers a way of automatically evaluating and eliminating ambiguity in pan-genomes.

**Author Summary:** Clustering of biological sequences is a critical step in studying bacterial species with large sequence diversity. Existing clustering approaches group sequences together based on similarity. However, these approaches do not offer a way of evaluating the confidence of their output. This makes it impossible to determine whether the clustering output reflect biologically relevant clusters. Most existing methods also do not allow cluster augmentation, which is the quick incorporation and clustering of newly available sequences with an already clustered set. We present Boundary-Forest Clustering (BFClust) as a method that can generate cluster confidence scores, as well as allow cluster augmentation. In addition to having these additional key functionalities and being scalable to large dataset sizes, BFClust matches and outperforms state-of-the-art software in terms of accuracy, robustness to noise and speed. We show on 4 *Streptococcus pneumoniae* datasets that the confidence scores uniquely generated by BFClust can indeed be used to identify ambiguous sequence clusters. These scores thereby serve as a quality control step before further analysis on the clustering output commences. BFClust is currently the only biological sequence clustering tool that allows augmentation and outputs confidence scores, which should benefit most pan-genome studies.

## Introduction

Most bacterial species harbor large amounts of sequence diversity. For example, any given strain of the human respiratory bacterial pathogen *Streptococcus pneumoniae* has about 2,100 genes in its genome, but two strains can differ by the presence or absence of hundreds of genes. In fact, the core genome (the genes shared by all strains) is estimated to be anywhere between 15-50% of the pangenome (the entire genetic repertoire of the species, thought to contain between 5,000-10,000 genes) (1–3). In species such as *S. pneumoniae* where there is a large amount of genetic diversity, phylogenetic studies or studies that compare multiple strains first necessitate identifying which genetic elements are the same across the different strains.

Establishing gene correspondence is often achieved by orthologue clustering, which groups orthologues of the same gene based on sequence similarity. An ideal orthologue clustering method is scalable, accurate, allows *cluster augmentation* (the addition of new sequences to a clustered set, without changing the initial clustering), and assigns a *confidence score* to the clusters it outputs. Earlier approaches for orthologue clustering such as PanOCT (4) and PGAP (5), involve all-against-all sequence comparisons, which compares each sequence to all other sequences in the dataset, and uses all of these comparisons to cluster. With such an approach, the number of comparisons increase quadratically with the number of data points, making these methods inapplicable for large datasets. Other approaches such as CD-HIT (6) and Usearch UCLUST (7) require the user to choose a sequence similarity threshold for the clusters. These *direct threshold methods* ensure that sequences that are more dissimilar than the threshold do not appear in the same cluster, and are extremely fast. CD-HIT has been used for pan-genome clustering for different microbial species (3,8,9), while UCLUST is the default clustering algorithm in the Bacterial Pan Genome Analysis tool (BPGA) (10), which is also used for multiple species’ pan-genome analysis (11–15). Importantly, when using *direct-threshold methods*, the correct value of the threshold may be difficult or impossible to determine, and an incorrectly chosen threshold value directly impacts clustering accuracy.

An alternative to *direct-threshold methods* are network-based methods, such as spectral clustering or Markov clustering (MCL) (16,17). These methods represent each sequence as a node in a network, and sequences are connected to one another according to how similar they are. The resulting network can then be partitioned into clusters based on its topology. Since generating the network requires all-against-all comparisons, these methods also do not scale out-of-the-box. To overcome this challenge, the two software solutions for pan-genome clustering, PanX (18) and Roary (19), first use a *representative selection step* – which reduces the redundancy in, and the size of, the dataset by grouping extremely similar sequences together. For each group, a representative sequence is picked, and the representatives are then clustered using MCL. The cluster membership for the representatives is then extrapolated to all sequences.

There are multiple strategies for *representative selection*. For instance, PanX separates consecutive input sequences into groups, then performs clustering within each one of these groups, and finally, selects one representative from each cluster from each group. Alternatively, Roary uses CD-HIT as its representative selection method (19). In either case, only a single set of representatives is selected, and there is no guarantee that this set *best* represents the whole dataset, which is a critical limitation.

Two additional challenges for pan-genome clustering are a lack of *cluster augmentation*, and a lack of *confidence scores* on the clustering output. Currently CD-HIT is the only clustering software that enables *cluster augmentation*, and no software produces *confidence scores*, which are critical in evaluating the ambiguity in the clustering results.

To overcome these challenges, we developed BFClust. BFClust uses a Boundary-Forest as a *representative selection* step, resulting in multiple sets of representatives that are stored. Each set of representatives is then clustered using MCL, yielding a *clustering ensemble*. A final *consensus clustering* step yields a single clustering solution from the *ensemble*. This approach has 2 main advantages: 1) multiple sets of representatives and *consensus clustering* enable calculation of *confidence scores*; 2) storing the Boundary-Forest enables quick *cluster augmentation*.

In this work, we evaluate the performance of 7 clustering methods (including hierarchical, *K*-means, spectral and MCL), and show that network-based methods such as MCL outperform others. BFClust using MCL is then compared to UCLUST, CD-HIT, PanX and Roary, which highlights that BFClust and PanX have high accuracy and robustness to noise when evaluated on a synthetic dataset generated *in silico* with known cluster assignments. In real pan-genome datasets, BFClust identifies clusters with low *confidence scores*, even in the core genome. Since such clusters most likely do not represent real orthologues, the *confidence score* can thus serve as a means to filter clustering results, only retaining unambiguous clusters. To the best of our knowledge, BFClust is the only clustering solution that produces the critical *confidence scores*, offers automatic *cluster augmentation*, and updates *confidence scores* during *cluster augmentation*. BFClust thereby is a unique method that enables quality control on the clusters it produces, while matching current gold standard tools in terms speed and performance.

## Results

### Algorithm Overview

Clustering of sequences using BFClust has three major steps: 1) *representative selection* i.e. reducing redundancy in the input data using Boundary-Forest; 2) clustering of each set of representatives associated with each Boundary-Tree into an ensemble of clustering solutions; and 3) deriving a *consensus clustering* from this ensemble of solutions (**Figure 1**). Once a *consensus clustering* is obtained, each cluster is assigned a *cluster confidence score*, and each amino acid sequence is given an *item consensus score*, based on the agreement of the clustering produced using the different Boundary-Trees.

**Figure 1:**
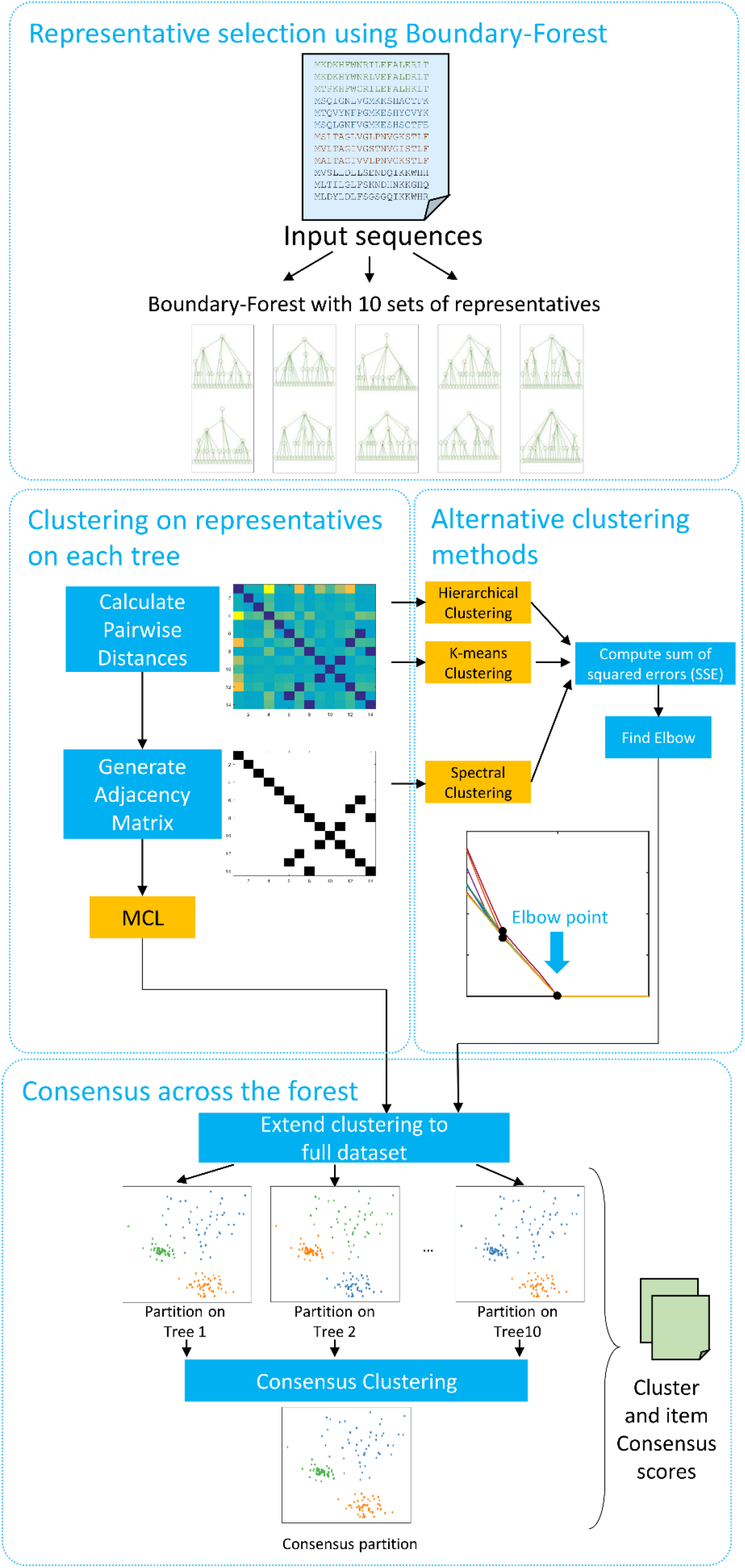
Algorithm overview. From the input sequences, multiple sets of representatives are selected using Boundary-Forest. Each set of representatives is stored as a Boundary-Tree. This reduces a large input dataset to a small set of representative sequences in the forest. Then, representatives on each tree are clustered using MCL. For comparison purposes, the following alternative algorithms were tested: Hierarchical, 2 variants of K-means, and 3 variants of Spectral clustering. Once representative sequences on each tree are clustered, the cluster assignments are extended to the full input sequence set, producing a clustering ensemble i.e. one clustering output associated with each set of representatives. A consensus clustering step is then used to take the clustering ensemble across the trees and produce a single clustering solution, as well as confidence scores. Cluster consensus scores are calculated for each cluster, and item consensus scores are calculated for each sequence within each cluster.

A naïve way to cluster all sequences from many bacterial genomes would be to look at all-vs-all pairwise sequence comparisons. Since all-vs-all pairwise comparisons require a computational effort that scales quadratically (O(*N*^2^) comparisons) with the number of sequences (*N*), it is beneficial to apply a *representative selection* scheme such that a group of extremely similar sequences is represented by a single sequence. We achieve this by constructing a Boundary-Forest (see **Supplementary Notes** for pseudocode). In a Boundary-Forest, *n* Boundary-Trees are constructed, with *n* =10 as the default size of the forest. Before constructing each Boundary-Tree, the order of sequences is randomized. The Boundary-Tree is constructed by placing the first sequence as the root, and the second sequence as its child. Then, each subsequent sequence is compared to the root node and its children. If the Smith-Waterman distance (20) between the incoming sequence and a node in the Boundary-Tree is smaller than a pre-set threshold *t*, the incoming sequence is represented by this node, and omitted from the tree. If the incoming sequence is not within the threshold of the root node or its children, we select the node (among the parent and children being compared to the incoming sequence) with smallest distance to the incoming sequence. If the newly selected node also has children, we repeat the comparison, moving down the tree until a representative is found that is sufficiently close to the incoming sequence. If such a node is found, we assign this node as the representative of the incoming sequence, and start processing the next incoming sequence. If no node within distance *t* is found on the tree, the new sequence is added as a child of a leaf in the tree. To control the breadth of the tree, we also limit the maximum number of children allowed for each node (with the parameter *“MaxChild”*). Note that below, we explore the sensitivity of the approach to these parameters. Since any Boundary-Tree that is constructed is sensitive to the order in which the sequences are read, a single tree is not guaranteed to capture a set of representatives that leads to highly accurate downstream clustering. Therefore, multiple Boundary-Trees (the Boundary-Forest) are used, which can be thought of as multiple ‘opinions’ on what representative sequences should be chosen. Once the sequence set is reduced to *n* sets of representatives, stored as a forest of *n* trees, the pairwise distances are computed within each set of representatives, and well-established clustering algorithms are applied.

After clustering the representatives, the cluster assignments of the representatives are *extended* to the full dataset. This is a necessary step for comparing the clustering output to the ground truth, comparing two clustering outputs to each other, and for *consensus clustering*, as these actions are performed on the full dataset, and not on the representatives. During the construction of each Boundary-Tree, each sequence is assigned a representative (or is itself a representative) based on sequence similarity. *Cluster extension* from the representatives to the full dataset is done by assigning each sequence the cluster of its representative.

The representatives of each Boundary-Tree are used to produce one clustering output, the whole Boundary-Forest thus leading to an *ensemble* of possible clustering outputs. *Consensus clustering* across the *clustering ensemble* is then applied, combining the clustering output obtained from each tree, to improve accuracy. Finally, BFClust compares how the different clustering outputs in the ensemble contribute to the *consensus clustering*, and using the differences in these contributions it assigns an *item confidence score* to the membership of each sequence to its consensus cluster, and a *cluster confidence score* to the existence of each cluster.

### Boundary-Forest reduces redundancy in the sequence set

In order to evaluate whether Boundary-Forest effectively reduces an input dataset into a small set of representatives by removing redundant sequences, we studied how much this step reduces the size of the dataset, how this reduction depends on the algorithm’s parameters, and how, in turn, this affects downstream clustering accuracy. We generated a small test dataset (‘minigenomes’), with 500 sequences of varying length (ranging from 65 to 1170 amino acids). This dataset has 50 noisy copies of 10 genes, and therefore 10 inherent sequence clusters. The noise is independent, random changes in the nucleotide sequence with probability 0.01 per nucleotide. Since BFClust uses amino acid sequences by default, the perturbed nucleotide sequences were then translated into perturbed amino acid sequences *in silico*. As the changes introduced to the sequences are random, 10 replicate sequence sets with the same mutation probability were generated. **Figure 2A** shows how the size of the Boundary-Tree constructed from this dataset is robust to two parameters that are crucial in constructing the Boundary-Tree: *MaxChild* and the sequence similarity threshold *t*. A detailed description of all parameters used in BFClust is provided in **S1 Appendix**. While a drastically small threshold value (*t* = 0.01) results in larger trees (which is intuitive, since with a smaller similarity threshold, fewer sequences can be represented by the same node), the size of the tree is robust to a large range of *t* and *MaxChild* values. Once a tree is generated, applying downstream clustering still may require all pairwise comparisons on the representatives. However, the number of pairwise comparisons are now greatly reduced; for example, a tree generated with *MaxChild* = 10 and *t* = 0.1 has 15 nodes (**Figure 2A**), which requires only 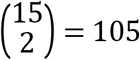 pairwise comparisons for clustering, versus 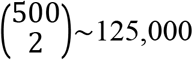 when clustering the entire dataset. Importantly, the construction of each Boundary-Tree also requires relatively few sequence comparisons itself: for example, 4500 comparisons are sufficient to generate the Boundary-Tree, when *MaxChild* > 2 (**Figure 2B**). The trees constructed are also relatively shallow (**Figure 2C**), meaning that the addition of a new sequence to an existing forest will require very few sequence comparisons (the number of comparisons grows with the depth of the tree) and will thus be very fast, which is relevant when we later discuss *clustering augmentation*. This is in line with the results reported by the creators of Boundary-Forest, where the depth of Boundary-Trees was shown to depend logarithmically on the number of data points for multiple datasets (21). Importantly, applying the full BFClust pipeline with varying the parameters *MaxChild* and *t* did not alter the clustering output, and the recovered clusters using any combination of these parameters (in the ranges presented in **Figure 2**) were identical to the ground truth and resulted in no error in clustering output.

**Figure 2:**
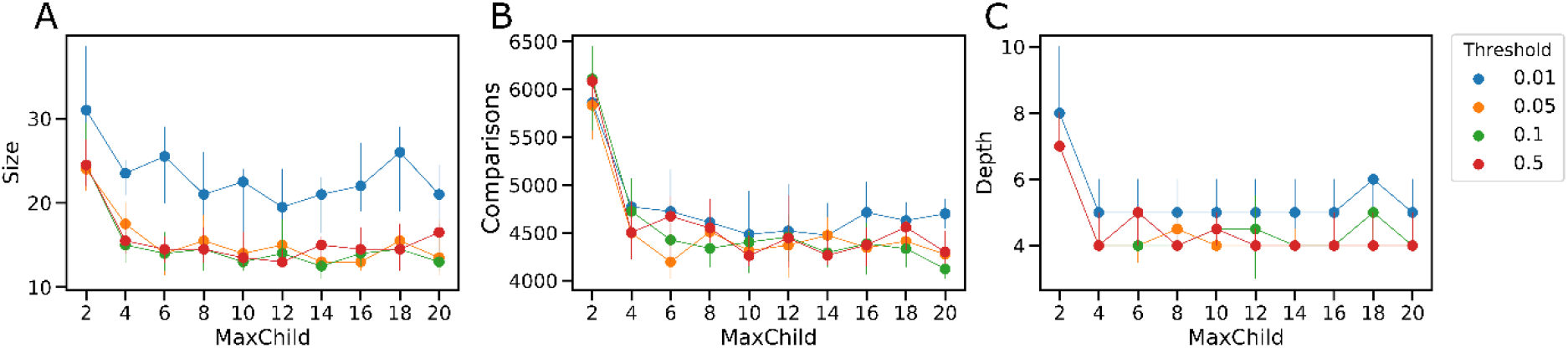
Boundary-Forest reduces redundancy in the sequence set. Boundary-Trees were generated from a 500-sequence dataset, in order to select representatives. The trees are small, shallow and quickly constructed. MaxChild: maximum number of children allowed for one node. Threshold: sequence similarity threshold, below which a sequence is assigned the tree node as a representative. **A.** The size (number of nodes) of the Boundary-Tree **B.** Number of calls made to the sequence comparison function **C.** The depth of the resulting tree, dependent on MaxChild and Threshold. Overall, the tree depth/size/number of calls made to construct the tree are robust to user defined parameters MaxChild and threshold. Points are the mean ± standard deviation for 10 replicates.

There are alternatives to *representative selection* that involve simpler algorithms than Boundary-Forest; two approaches we discuss here are a *random sampling* and a *naïve sampling* strategy. *Random sampling* is the selection of representatives randomly from the full set of sequences. This is not a viable strategy, as it does not guarantee that all clusters will be selected. For example, on the set of 500 sequences with 10 known clusters, a random selection is likely to include at least one representative from all 10 clusters only when a sufficiently large number of sequences (e.g. *N* = 50) are randomly selected (**Supplementary Figure 1**). In this case, a 10-fold reduction in the number of representatives (compared to the full sequence set) may seem promising. However, in real pan-genome datasets, this reduction might result in hundreds of thousands of representative sequences, which would still be prohibitively numerous for downstream clustering. Moreover, in real sequence sets it is not clear how many sequence clusters are present, and *random sampling* risks missing smaller clusters. Thus, estimating the number of random samples to be selected such that all clusters will be represented is difficult, if not impossible.

In the second, *naïve sampling* scheme, sequences are read in a random order, and the first sequence is placed into a ‘representatives’ group. Each incoming sequence is then compared to the existing representatives, and if no representative closer than a threshold *t* is found, the incoming sequence is added to the representatives group. Both CD-HIT and UCLUST apply this *naïve sampling* strategy. On the small 500-sequence minigenomes dataset, the number of representatives selected by the *naïve sampling* and Boundary-Tree are similar (**Table 1**). However, *cluster augmentation*, which is a key advance we present below, requires the comparison of new sequences to all of the representatives in the case of *naïve sampling*, while in the Boundary-Tree, the traversal of a much smaller subset of representatives is required. The number of comparisons on a Boundary-Tree is dependent on the tree depth, and the number of children each node has, which is limited with the *MaxChild* parameter. With *MaxChild* = 10, the estimated number of possible comparisons in the Boundary-Tree for new sequences would be at worst *10* × (*tree depth*), whereas in the naïve scheme it would be equal to the current size of the representatives set. The advantage of the Boundary-Forest becomes apparent when a larger sequence set is considered. For instance, 20 *S. pneumoniae* strains were selected from the RefSeq database (**Supplementary Table 1**), and the coding sequences were subjected to *naïve sampling* and Boundary-Tree sampling. While the number of representatives in the Boundary-Tree is about twice as large as the representatives picked with naïve sampling, the trees are shallow. The number of comparisons needed to process a new sequence in the Boundary-Tree is ~90, which is about 35-fold smaller than the comparisons using the naïve representative set (~3265). Therefore, we conclude that the extra effort at the beginning of constructing the Boundary-Forest results in more efficient sample processing as the sequence dataset grows larger.

**Table 1:** Comparison of *naïve sampling* and Boundary-Trees as *representative selection* methods. Representatives were selected from two datasets (minigenomes, a synthetic small sequence set; and RefSeq, sequences from 20 *S. pneumoniae* strains present in the RefSeq database), using either *naïve sampling* or Boundary-Trees. N: number of sequences in the dataset. Representatives: number of representatives selected with *naïve sampling* or Boundary-Tree. Tree depth: depth of a Boundary-Tree. Mean±standard deviation of 10 replicate sets of representatives are reported. BT comparisons: expected number of comparisons that will be made during *cluster augmentation* using Boundary-Trees (note this value is the same as tree depth multiplied by the number of children allowed, which is 10 by default).

| Dataset | N | Representatives<br>(Naïve Sampling) | Representatives<br>(Boundary-Tree) | Tree depth | BT<br>comparisons |
| --- | --- | --- | --- | --- | --- |
| minigenomes | 500 | 15.5±5.6 | 13.2±2.0 | 4.5±0.8 | 45 |
| RefSeq | 42010 | 3264.7±5.4 | 6579.8±78.3 | 9±0.82 | 90 |

### MCL outperforms other methods during clustering of the representative sequences

We tested 4 commonly used clustering methods (a total of 7 variants of the 4 methods) for downstream clustering of the representative sequences obtained from the Boundary-Forest. These are 1) hierarchical clustering with ward linkage (‘Ward’), 2) *K*-means clustering of sequence distances (‘Kmeans’), 3) *K*-means clustering on vector-embeddings of amino acid sequences (‘Kmeans V’), 4) non-normalized spectral clustering (‘Spectral NN’), 5) spectral clustering with Shi-Malik normalization (‘Spectral SM’) (22), 6) spectral clustering with Jordan-Weiss-Ng normalization (‘Spectral JWN’) (23), and 7) Markov clustering (MCL) (24). With the exception of MCL, these methods do not automatically select the number of clusters to output, and rely on the user to specify this value. Given a set of biological sequences, it is not inherently evident how many sequence clusters there should be. To address this, each method is applied multiple times, each time generating partitions of the data with a different number of clusters. Then, a curve of the sum of squared errors (SSE) as a function of different cluster sizes is computed. SSE measures the scatter within each cluster and thus can be used as a quality metric (25). Finally, the most appropriate number of clusters, *K* is determined by finding the ‘elbow’ on the SSE curve. The elbow point is defined as the point where the second derivative of the SSE curve is maximized (see Methods, and **Supplementary Figure 2**). Intuitively, the elbow point is the simplest model (i.e. smallest *K*) beyond which adding more complexity (increasing *K*) results in substantially lower gains in terms of SSE i.e. error is lowered much slower after the elbow point. Using this procedure, for each method, applied on representatives on each tree, the SSE curve is computed, and the elbow points are selected (**Figure 3A**, black points). As expected, with increasing number of clusters the total SSE decreases (**Figure 3A**). In the example in this figure, where the input sequences have 10 clusters, SSE is minimized at 0 for number of clusters *K* ≥ 10. In real genomic datasets, it might not be the case that SSE is minimized to 0 when the correct *K* is selected. Therefore, the elbow method described above is preferable over selecting the *K* at which the SSE curve attains its minimum.

**Figure 3:**
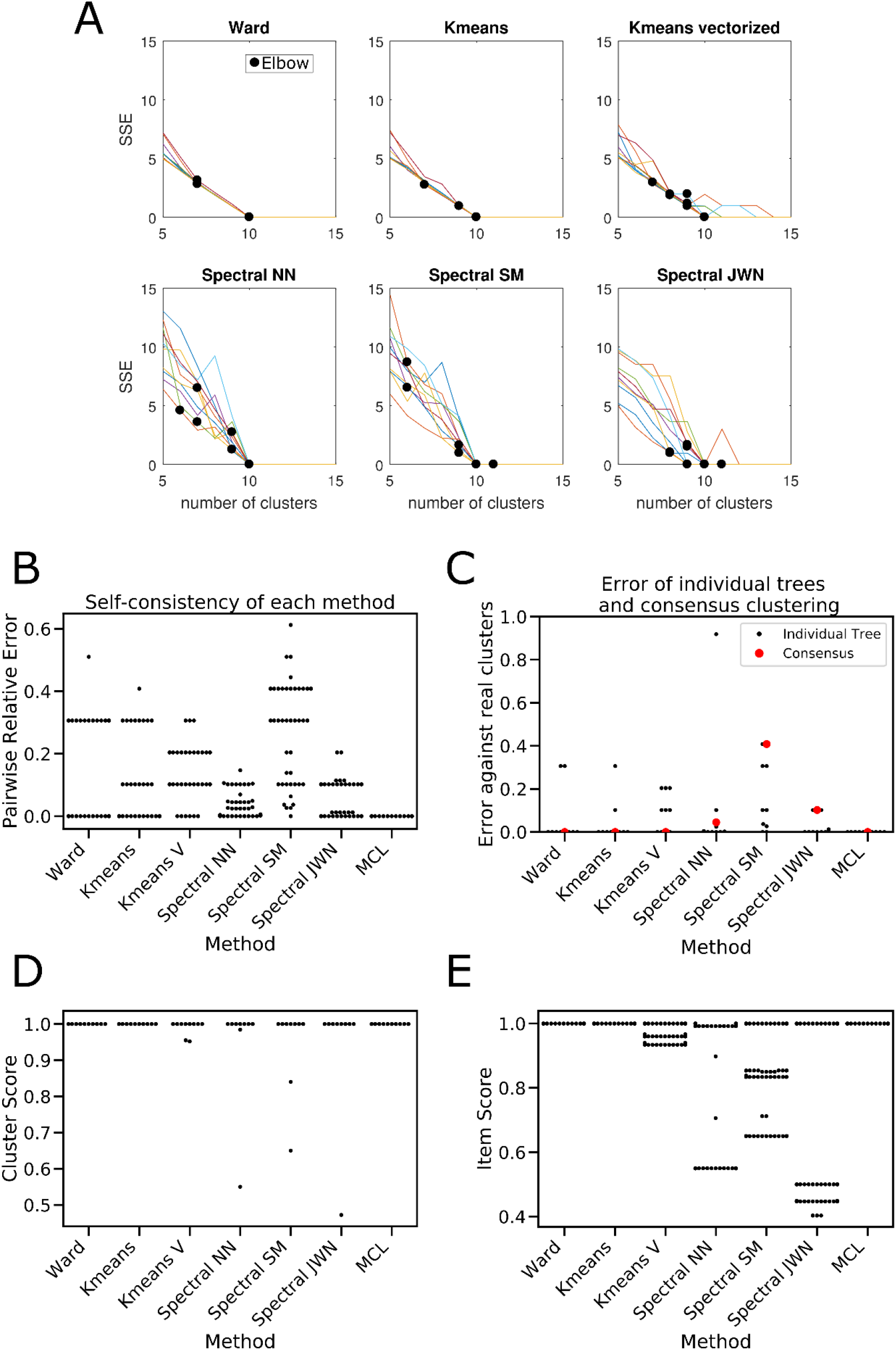
Evaluation of downstream methods of clustering on the reduced data. **A.** Sum of squared errors (SSE) is the metric used to detect the number of clusters for methods other than MCL. Each trace is a clustering output applied to representatives on a different Boundary-Tree, scanning increasing number of clusters. Black dots: automatically detected elbow points on each trace (note that these elbow points are often overlapping). **B.** Self-consistency of each downstream method. The clustering output for the elbow points were compared across 10 different trees for the same method. **C.** Error against real cluster assignments of each individual tree (black dots) and the consensus (red squares) among the forest, for each method. Relative error: error divided by maximum possible error. **D.** Cluster confidence score for each recovered cluster (*n* = 10 clusters). **E.** Item confidence score for each clustered sequence (with *n*=500 sequences)

The selected elbow on the SSE curve for each tree is often at the correct *K*, but not always (**Figure 3A**). This is because the multiple Boundary-Trees are different, and their construction depends on the order in which the sequences are processed. Therefore, the same clustering method can result in different clustering partitions, depending on the Boundary-Tree used to reduce the data. If we compare the amount of disagreement among the trees for the same method, we see that different trees do indeed result in some inconsistencies (**Figure 3B**). Notably, MCL is the one downstream method that is extremely self-consistent. To reconcile the remaining inconsistencies, which improves the accuracy of the clustering, and to be able to report *confidence scores*, we add a *consensus clustering* step to BFClust. For this, we take the clustering assignments from the clustering obtained from each tree as a vector, and use a scalable clustering method (e.g. *K*-medoids) to cluster these vectors (**Supplementary Figure 3**). The median of the elbow *K* values from each tree is used as the final number of clusters to be generated by the consensus. **Figure 3C** shows that after the consensus step, the errors against the known cluster assignments are reduced to 0 for most methods, despite the clustering from individual trees having higher error prior to consensus.

### BFClust can compute cluster confidence scores

The consensus clustering step across the Boundary-Forest not only reduces error, but it also allows confidence estimation for the existence of each cluster, and for the membership of each sequence in its consensus cluster. By comparing the clustering done using the representatives on each Boundary-Tree, it is possible to measure how frequently a cluster has the same members, and use this value as an estimate of cluster confidence. We define a *cluster confidence score* (for each cluster) and an *item confidence score* (for each sequence), and include both sets of values in the BFClust output. Both values depend on the *consensus index* (26). The *consensus index* of a pair of sequences *i* and *j* is the number of times that they appear together in the same cluster across *n* Boundary-Trees, divided by the total number *n* of Boundary-Trees used. The *item confidence score* for item *i* is the average *consensus index* between *i* and all other members of the same consensus cluster. The *cluster confidence score* is the average *consensus index* between all pairs of items within the same consensus cluster (**Supplementary Figure 3**). Both scores take a value between 0 and 1, and a score of 1 indicates perfect agreement of cluster memberships across the Boundary-Forest. The *cluster confidence score* (**Figure 3D**) and the *item confidence score* (**Figure 3E**) was computed on the minigenomes dataset with a mutation rate = 0.1. Spectral clustering variants result in a few clusters with low *confidence scores*, whereas Ward and MCL clustering have the highest scores, indicating these methods have high agreement across the Boundary-Forest, and produce stable clusters even in the presence of noise.

In **Figures 3D** and **3E**, the cluster and item *confidence scores* are presented as a means to evaluate the self-consistency of each method. However, when clustering real pan-genome datasets, these *confidence scores* computed for each cluster or sequence serve as a confidence measure of the clustering output. This means that clusters with lower scores are those with higher uncertainty, and should be used with caution in further analysis and interpretation.

### Cluster augmentation: addition of new sequences to an existing clustering

A major advantage of the BFClust algorithm is that it stores the Boundary-Forest containing representatives from all previously processed input sequences. This allows BFClust to add new sequences to an existing clustering/partition while being able to update the confidence scores without much computational work. To achieve this, we implement a *cluster augmentation* method (see **Figure 4A** for a schematic overview). A set of incoming input sequences can either be used as-is, or optionally are reduced to a new set of representatives by constructing a new Boundary-Tree. These new sequences (or representatives) are then run through each tree in the existing forest (corresponding to the already clustered set of sequences), and for each new representative, a close match on each of the 10 trees is identified. The cluster membership associated with each tree is extracted for these close matches from the previously computed clustering. Each new sequence is assigned the same clustering membership as that of the corresponding close match within each tree. This results in a vector of cluster assignments for each new sequence. Afterwards, the vectors composed of the cluster memberships for the new input representatives from each tree are turned into a consensus cluster assignment using a nearest neighbor search on the cluster assignments of the datapoints in the existing dataset. If an initial *representative selection* step was used, the *consensus clustering* on each input representative is then extended to the entire input set, using the same procedure of *cluster extension* during *de novo* clustering.

**Figure 4:**
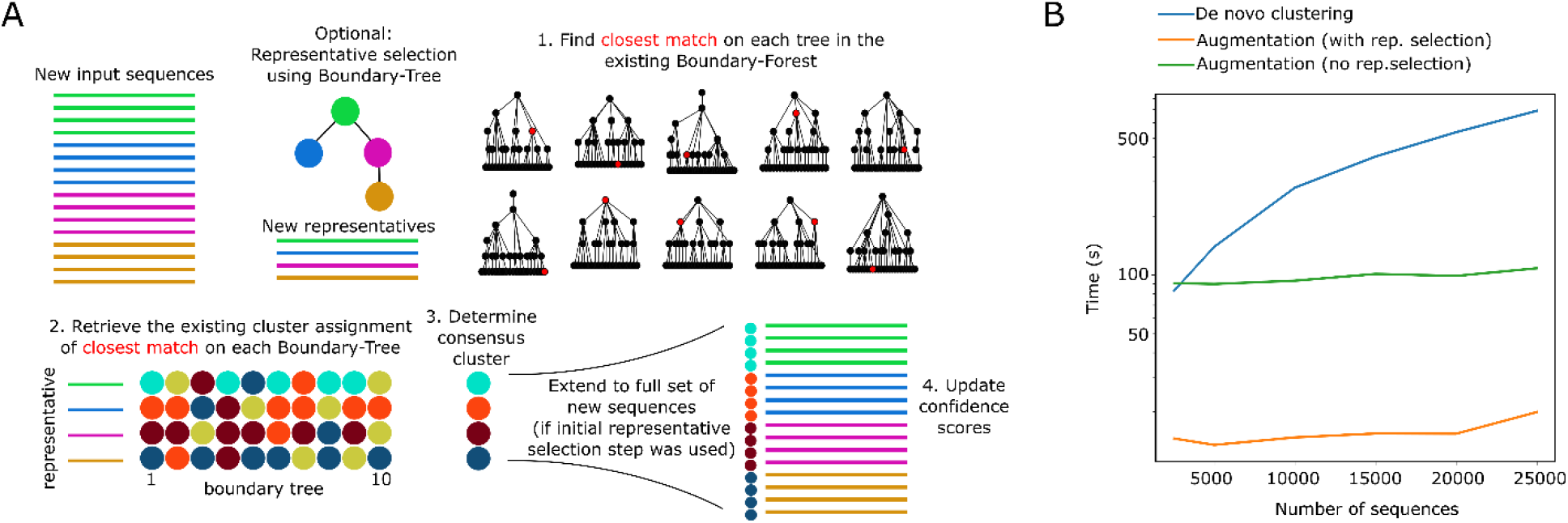
BFClust allows cluster augmentation. **A.** Cluster augmentation method overview/schematic. The incoming sequences are either processed as-is, or they can be reduced to a small set of representatives using a Boundary-Tree. The new sequences (or representatives) are compared to the existing Boundary-Forest associated with the already clustered dataset. A close match in each tree, for each input sequence is found (red nodes). The cluster assignments of these closest matches are retrieved, and a consensus cluster assignment is computed using a nearest neighbor search. If representative selection is used, the consensus clusters assigned to the new representatives are extended to the full input dataset. The cluster assignments of the new sequences, as well as updated confidence scores for both the existing and new sequences are produced as the output. **B.** Cluster augmentation is faster than clustering *de novo*. Runtime of clustering sequences *de novo* (blue), or cluster augmentation onto an already clustered set (orange, green). For augmentation, N-500 sequences were clustered *de novo*, and the runtime for the augmentation of the remaining 500 sequences is reported. The runtime with and without an optional representative selection step is shown as orange and green lines respectively. The additional representative selection step improves runtime further.

The runtime of *de novo* clustering, and *cluster augmentation* scales tractably with increasing number of data points (**Figure 4B**). For *de novo* clustering, increasing numbers of replicates of the 500-sequence dataset (with different random mutations) were included. For *cluster augmentation*, the last 500 sequences from the dataset was left out, the remaining *N* − 500 sequences were clustered *de novo*, and the remaining 500 sequences were added to the dataset using the procedure described above. Augmentation is much faster than clustering sequences *de novo*, even without the initial *representative selection* step (**Figure 4B**, green curve). When the optional *representative selection* step is included, there is an additional 5-fold increase in the runtime (**Figure 4B**, orange curve). Error on the final clustering, both for *de novo*, and for *cluster augmentation* was 0.

### Comparison and benchmarking with existing methods

We selected four existing sequence clustering methods to compare BFClust against. The first is Usearch UCLUST (7), a very fast and scalable algorithm that is also the basis of several other pan-genome phylogenetic analysis pipelines such as SaturnV (27), PanPhlAn (28) and BPGA (10). Second, we consider CD-HIT, another scalable software that has been used directly in pan-genome analysis (8,9) and for *representative selection* in other pipelines such as Roary (19). Roary is also included, as it was one of the first popular software tools that allowed the pan-genome analysis of several hundred genomes at once, and therefore has been utilized in recent studies (29,30). Finally, PanX (18) is included, another recent software that can be used with hundreds of genomes.

First, the runtime of the four software tools was compared to BFClust (**Figure 5A**). The *direct-threshold methods* UCLUST and CD-HIT are orders of magnitude faster than the other methods. On the other hand, methods that employ network-based clustering (MCL for BFClust, PanX and Roary) take far longer. With 25,000 input sequences, BFClust and Roary have similar runtimes, about 50x shorter than PanX (**Figure 5A, Table 2**).

**Figure 5:**
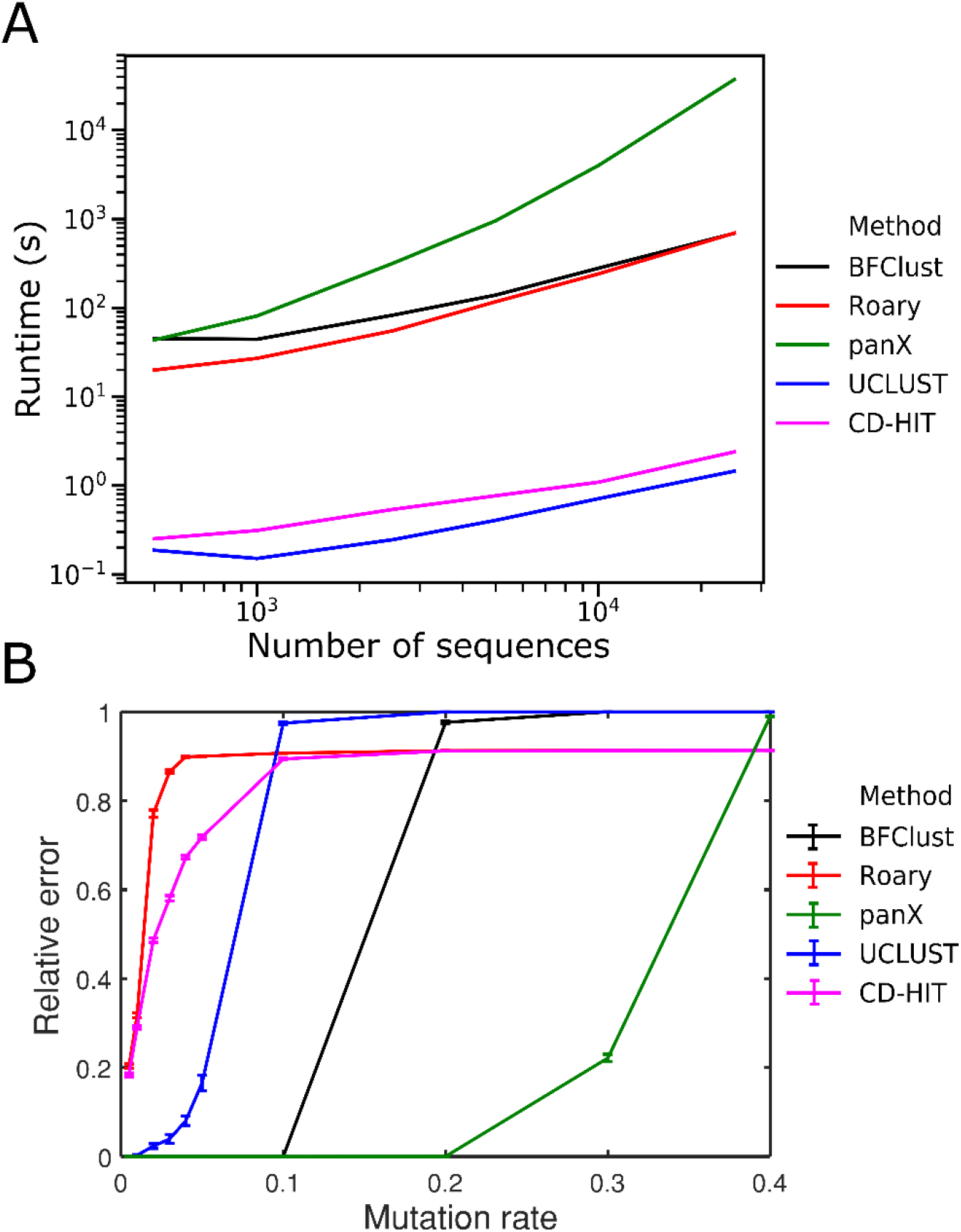
Comparison of BFClust to existing methods. **A.** Runtime in seconds of each method, as a function of dataset size (number of input sequences). **B.** Sensitivity to noise of each method. Relative error against known clusters increases for all methods with increasing amount of mutations in the data. BFClust and PanX have 0 error when mutation rate is ≤ 0.1. Mutation rate: the probability that each nucleotide is replaced by a random one. Mean ± standard error of 10 replicates are shown by the error bars.

**Table 2:** Comparison of software tools applied to pan-genome-wide orthologue clustering. Runtime: time it takes to run each method on 25,000 sequences (in seconds). Relative Error: Clustering error compared to ground truth on the minigenomes set with mutation rate = 0.1. Representative selection: whether the clustering strategy reduces the input dataset to a small set of representatives before/during clustering. Cluster augmentation: whether the method provides an automatic procedure for adding new sequences to an existing clustering partition. Confidence score: whether the clustering algorithm returns a clustering consensus score as an output. Network-based clustering: whether the method used network-based clustering strategies (MCL). ^b^: PanX uses a divide-and-conquer strategy to process large datasets, where batches of 50 genomes are clustered at one time. Representatives from each cluster are selected in each batch. Since representative selection is done after clustering a relatively large set of sequences, we consider this strategy substantially different than other representative selection strategies.

| Method | Runtime (s) | Relative Error | Reference | Representative selection | Cluster Augmentation | Confidence score | Network-based clustering |
| --- | --- | --- | --- | --- | --- | --- | --- |
| BFClust | 691.6 | 0 | This work | Yes | Yes | Yes | Yes |
| PanX | 37228.6 | 0 | (18) | No<sup>b</sup> | No | No | Yes |
| Roary | 695.0 | 0.907 | (19) | Yes | No | No | Yes |
| CD-HIT | 2.4 | 0.894 | (6) | Yes | Yes | No | No |
| USEARCH | 1.5 | 0.975 | (7) | Yes | No | No | No |

In order to compare the sensitivity to noise of our approach to existing methods for biological sequence clustering, we generated an extended set of test sequences. Here, we made 10 replicates of the 500-sequence minigenome set (with 10 known gene clusters) at given mutation rates, ranging from 0 to 0.4. We then computed the error against known clusters for each mutation rate. Both Roary and its *representative selection* method CD-HIT are very sensitive to low levels of noise in the sequence data (**Figure 5B**). In contrast, BFClust and PanX have 0 error when the mutation rate is less than or equal to 0.1 (**Figure 5B, Table 2**). Based on this, we recommend using PanX or BFClust when a high amount of variation is expected in the sequence data, either due to genetic variation, or noise from error-prone sequencing technologies such as Oxford Nanopore (31). Error of BFClust when using other clustering methods is also low at mutation rates ≤ 0.1, however none of these methods outperform MCL (**Supplementary Figure 4**). In addition to being noise-insensitive, BFClust provides additional *confidence* scores, which are critical when clustering data with high sequence variability. An overall comparison of all 5 approaches are summarized in **Table 2**. BFClust and CD-HIT have the added advantage of allowing *cluster augmentation*. Importantly, BFClust is unique in the way it can output *cluster* and *item confidence scores*. In the application to real pan-genome clustering, these scores can be taken as a measure of cluster robustness.

### Clustering of real pan-genomes

To demonstrate the applicability of BFClust beyond synthetic datasets, i.e. on real pan-genomes, we explored several *S. pneumoniae* datasets. *S. pneumoniae* is a naturally competent, opportunistic human pathogen that is known to have a relatively large pan-genome, partially shaped by recombination events (1,2). Since in a real pan-genome, the ground truth for clustering is unknown, it is not possible to compute clustering error. Therefore, in this section we compare different clustering methods to each other and see whether they yield consistent outputs, in addition to exploring the *cluster confidence scores* generated by BFClust. Previously, core and pan-genome analyses using Roary had revealed that across different datasets of pneumococcal isolates, the core genome is not conserved, and the size of the pan-genome is not the same across datasets (3). However, it is unclear whether this is a consequence of the datasets (which come from different types of populations that are also geographically separated) being different from one another, or an artifact of the clustering method used. In order to avoid any bias associated with a specific dataset, we compiled 4 datasets in this study: 1) RefSeq (closed, chromosomal genomes, n=20) (32) 2) Maela (annotated contigs from a Thai refugee camp, n=348)(33); 3) Nijmegen (annotated contigs from a Dutch hospital, n=350)(34); and 4) MA (annotated contigs from surveillance data from Massachusetts, n=616)(2). Despite being the smallest dataset, the RefSeq set is the most diverse, as these strains have collection dates and countries of origin that vary the most (see **Supplementary Table 1** for a full list of strains). The Nijmegen dataset is comprised of pneumococcal isolates from invasive pneumococcal disease patients, whereas the MA and Maela datasets are collections of pneumococcal isolates from healthy individuals (i.e. carriage isolates).

We evaluated whether the core and accessory genome profiles detected by each method are consistent. A reasonable expectation for a given tool is that it produces similar core and pan-genome size estimates for the 3 larger datasets (MA, Maela, Nijmegen). This expectation is met by all methods but Roary, which shows a big discrepancy in the core and pan genome size across these datasets (**Figure 6A, B**). Relative to the other methods, it appears that Roary underestimates the core genome size, and over-estimates the pan-genome size (**Figure 6A, B**). In comparison, BFClust and PanX both find a larger core genome and a smaller accessory genome compared to the other methods, whereas UCLUST and CD-HIT find a similarly sized core genome, but a larger accessory genome compared to BFClust and PanX (**Figure 6**).

**Figure 6:**
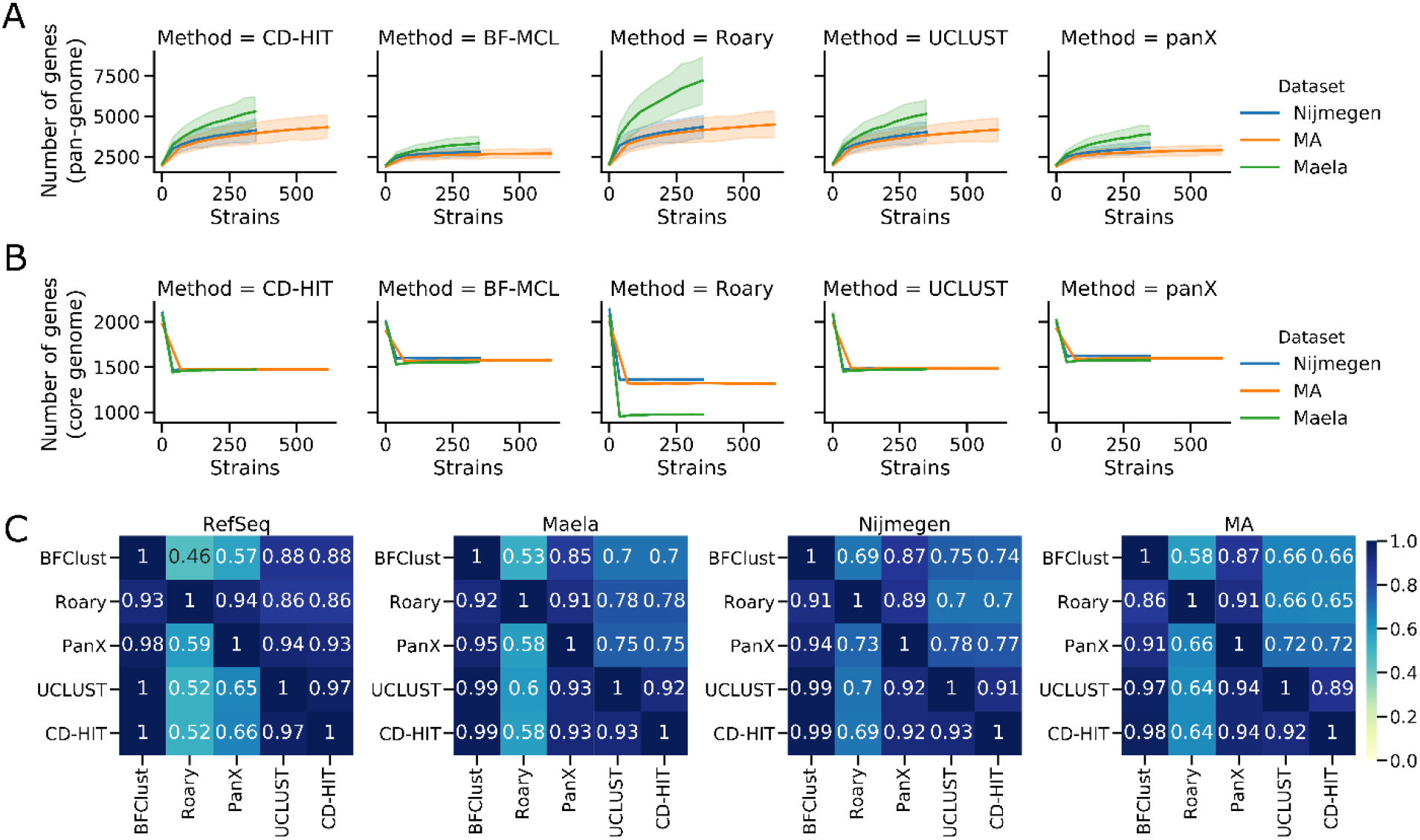
Clustering of real pan-genome sequences reveals differences across methods as well as datasets. **A.** Pan-genome size (total number of genes in the pan-genome) as a function of number of strains considered. **B.** Core genome size (total number of genes common across strains) as a function of strains considered. **C.** Cluster overlap (see methods) between different methods for each dataset. For **A.** and **B.** mean ± standard error of 10 replicates are shown by the line and error bands.

In these analyses, BFClust is used with MCL. The alternative method were also tested within the BFClust pipeline, on each of the 4 datasets. The non-network methods Ward, Kmeans and Kmeans-V do not find elbows on the SSE curves (**Supplementary Figure 5**), indicating that these methods are not finding meaningful clusters in these datasets. In contrast, all 3 variants of network-based Spectral clustering can find a clear elbow, and therefore may be more suitable for clustering of biological sequences

In order to compare the agreement between clustering methods on a given dataset, we computed cluster overlap: the proportion of clusters generated by one method that are fully contained within another cluster generated by a second method (note that this measure is sensitive to the direction of comparison; agreement of method A with B is not necessarily the same as the agreement of B with A; see Methods). Interestingly, on the same datasets, CD-HIT and UCLUST had the highest agreement, as determined by cluster overlap (**Figure 6C**). BFClust and PanX were also in high agreement. Roary appears to have poor agreement with other methods in one direction, which could be attributed to the fact that it is producing too many clusters, fewer of which end up in the core genome. This is potentially a consequence of Roary using CD-HIT for the first step of selecting representative sequences, as both were sensitive to noise.

Clustering of pan-genome sequences can be the first step of phylogenetic analyses. For instance, the SNPs within the core genome can be used to generate phylogenetic trees and make conclusions on population structure (2). In these analyses, it is essential that the clustering is unambiguous; incorrect clustering would potentially lead to misleading conclusions. We therefore computed the *cluster confidence scores* for each cluster obtained using BFClust, on each of the 4 *S. pneumoniae* datasets. The majority of the clusters had a score near 1, indicating very little ambiguity in the clustering output (**Figure 7**). Specifically, we observe high-confidence clustering in the core genome; the mean score for core genome clusters is > 0.999 (and the median score = 1) in all 4 datasets. In the 3 larger datasets (Maela, MA and Nijmegen), we observe that the lower scoring clusters are mainly in the accessory genome, shared by less than a third of the strains included. In all datasets, there exists a single cluster with a much lower score than the average, present in the majority (and in some datasets all) of the strains included (marked in red in **Figure 7**). This cluster is comprised mostly of sequences of very short length (~30 amino acids), annotated as hypothetical genes. It is unclear whether these short sequences are artifacts of sequencing errors, annotation errors, incomplete genome assembly or a combination of these factors. In any case, exclusion of such low-scoring clusters from downstream pan-genome/phylogenetic analyses would potentially increase confidence in those results. By providing a *cluster confidence score*, BFClust allows for screening clusters, and including only high-confidence ones in downstream analyses.

**Figure 7:**
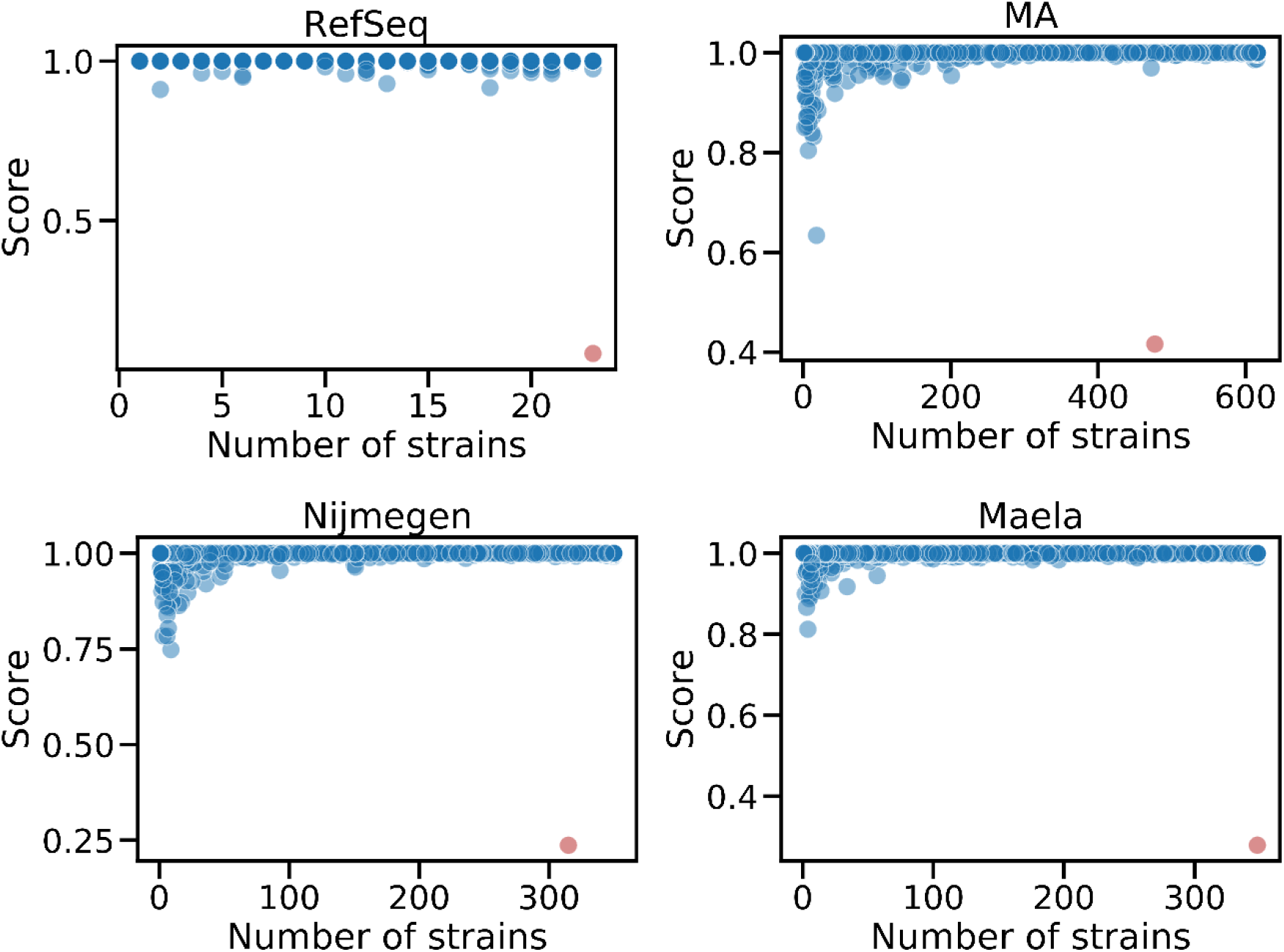
Cluster confidence scores for each cluster found using BFClust for 4 *S. pneumoniae* datasets, plotted against the number of strains that share the cluster. In general, the clusters with lower scores appear in the accessory genomes, and are not shared by many strains. There is one cluster within the core genome of each dataset with a low score (red clusters).

## Discussion

When clustering a set of sequences from a bacterial pan-genome, there are multiple options regarding the software/algorithm to choose from. We observed that *direct-threshold methods*, which are extremely fast (UCLUST and CD-HIT), have the advantage of scalability, but they often do poorly in terms of accuracy and sensitivity to noise (**Figure 5**). They also require the user to select a sequence similarity threshold, assuming all sequence clusters have similar sequence diversity, which is not always true. Different genetic elements are subject to different selective pressures, and therefore sequence conservation/diversification may be associated with multiple factors, e.g. essentiality (35), rendering the use of a single threshold problematic. Therefore, it is more advisable to first reduce the dataset into a smaller representative set (potentially using these faster methods) and then applying a more rigorous clustering method.

For the selection of representative sequences, we propose the use of Boundary-Forest, which is supported by existing numerical experiments showing the improved accuracy and speed of Boundary-Forest compared to other algorithms (21). Its implementation is also very simple (see pseudocode in **Supplementary Appendix**). The inclusion of multiple trees in the forest, and downstream application of *consensus clustering* also reduces errors, and results in BFClust being highly tolerant to noise, especially when BFClust is used with network-based downstream methods such as MCL. Furthermore, the use of multiple Boundary-Trees makes it possible to compute *confidence scores*. Saving a copy of the shallow Boundary-Trees allows rapid *cluster augmentation*, without having to alter the existing clustering assignments, which is highly desirable for consistency. Moreover, *augmentation* can be done while updating the clustering *confidence scores*. This makes BFClust the only pan-genome clustering method that can generate such a *cluster confidence score*, both during *de novo* clustering and during *cluster augmentation*.

The *cluster augmentation* strategy implemented in BFClust (and in CD-HIT) is distinct from online clustering methods. An online method updates the clustering, as new data points become available. This can potentially change the cluster memberships of the already-clustered dataset. BFClust on the other hand performs *cluster augmentation* by using a *K*-nearest neighbor search to find a cluster in the existing dataset that is a close match of the incoming sequence, without altering the existing clustering. This *K*-nearest neighbors search could potentially be replaced by a *K*-means or *K*-medioids clustering on the full set of already-clustered and incoming sequences together, in order to make BFClust more similar to an online method. However, it is not clear whether the same *K* value (total number of clusters, which was used in the initial clustering) would apply to the full set with the incoming sequences included. BIRCH (36) and stream clustering (37) are two examples of online clustering algorithms, however it is not likely that they would apply well to biological sequences, as they are online versions of hierarchical and *K*-means clustering respectively. We have shown in this work that these non-network-based algorithms fail to find a clear elbow point on real pan-genome sequence sets when clustering (**Supplementary Figure 5**). This suggests that non-network-based methods perform well in a simple case, where clusters are generated synthetically by accumulating random mutations; whereas real sequences are subject to different selective pressures and may not diverge from each other as uniformly as in the synthetic case. We therefore advocate the use of network-based methods, specifically MCL, for clustering of biological sequences.

The BFClust strategy has a number of advantageous features that can be explored further. Since each of the trees generated in BFClust has a small depth, the number of comparisons one needs to make for a new sequence set is relatively small (*tree depth* × 10 trees). Thus, this method offers a framework that makes the rapid integration of new clinically important isolates and their sequences possible. In the same vein, it is possible to quickly compare the clustering results of two different datasets (e.g. isolates of the same species of bacteria, collected from different geographical locations) by running one set through the Boundary-Forest of the other. Moreover, the networks that are generated as intermediate steps in clustering may harbor novel data that remains unexplored in this work. For instance, it may be possible to extract additional information from the network connectivity of sequences, and infer evolutionary trajectories of different genes under differing selective pressures (38).

In conclusion, UCLUST and CD-HIT may not be best suited for pan-genome clustering, as they depend on a user-supplied similarity threshold. CD-HIT and Roary (which uses CD-HIT) are especially sensitive to noise in the data. Nevertheless, the speed of UCLUST and CD-HIT make these methods attractive alternatives to BLAST when querying large datasets. Overall, BFClust and PanX are pan-genome orthologue clustering methods that are in high agreement, and can tolerate noise in the sequence dataset, although PanX has a higher runtime than BFClust. PanX has the advantage of informative and interactive visualizations, whereas BFClust has the added features of estimating *confidence scores*. Moreover, most pan-genome clustering methods (with the exception of CD-HIT) do not readily allow *cluster augmentation*, and to the best of our knowledge, no previous clustering method enables *cluster augmentation* while being able to output *confidence scores*. *Confidence scores* are crucial in pan-genome clustering, as they allow the researcher to avoid using ambiguous clusters (i.e. clusters with a low score) in downstream analyses and interpretation. With the *confidence score* of BFClust, such clusters can automatically be detected and excluded from further analysis.

## Materials and Methods

### Minigenome sequence sets

Nucleotide sequences spanning the first 10 annotated CDS sequences from S. pneumoniae strain TIGR4 were selected (nucleotides 1-27310, spanning locus tags: SP0001-SP0010) and used as an initial synthetic “minigenome”. Each of the minigenomes dataset contains 50 copies of these 10 genes, where random independent nucleotide mutations are allowed at a rate *r*. The mutation rate *r* is equal to the probability that one nucleotide is replaced with a different random nucleotide. We generated 100 such nucleotide-based “minigenomes” datasets, namely, 10 datasets for each of 10 different values of *r*, ranging from *r* = 0 to *r* = 0.4. As BFClust uses amino acid sequences by default, the nucleotide sequences for each gene were translated into amino acid sequences. To use Roary and panX, the nucleotide sequences and CDS annotations were converted into GFF3 and genbank files respectively.

### *Streptococcus pneumoniae* datasets

The full list of isolates used for clustering can be found in (**Supplementary Table 1**). The “RefSeq” dataset (*N* = 23) contains 21 annotated chromosome sequences from the RefSeq database (O’Leary et al, 2016) and 2 strains our lab uses in its studies: BHN97 (39) and 22F-CT (CDC Pneumococcal surveillance isolate). The “MA” dataset (*N* = 616) is a set of isolates from (2), collected from children between 2000-2007 from Massachusetts. The “Nijmegen” dataset (*N* = 350) includes isolates from invasive pneumococcal disease (IPD) patients in Nijmegen, Netherlands, collected between 2001-2011 (34). The “Maela” dataset (*N* = 348) include a random subset of carriage isolates collected from the Maela refugee camp in Thailand between 2007-2010 (33).

For strains BHN97 and 22F-CT, and the Nijmegen and MA datasets, unannotated contigs were assembled into closed, circular chromosomes, and annotated using the PATRIC-RAST annotation server (40). The genomes were then converted to genbank format, with *dnaA* as the first coding sequence using custom in-house scripts. The translated sequences of all CDS annotations were then extracted into a fasta file for each dataset using Biopython (41). When necessary, the genbank files were converted to GFF3 for use with Roary.

### Boundary-Forest

Within BFClust, a large sequence dataset is reduced to a set of representative sequences using Boundary-Forests. For each input dataset, 10 randomized read orders are generated. The sequences are read in these orders and 10 different Boundary-Trees are constructed as described in (21). Briefly, the first sequence that is read is placed as the root node, and the second as its child. For each subsequent sequence read, it is compared to the root node, and all its children using Smith-Waterman distance (20). If the sequence being processed is within a pre-determined distance similarity threshold *t* of a node already on the tree, then this node on the tree becomes its representative. This means that the sequence being processed is marked with the identity of the representative, and is not included in the tree. Otherwise, the sequence is compared to the current node, and all its children, and added to the tree as a child of the node that it is closest to. Most of the input sequences are not included in the tree and are simply associated with a representative on the tree. Boundary-Trees contain ~2% of the original input sequences, making the clustering of large numbers of (e.g. ~1 million) sequences possible. By default, the sequence distance similarity threshold is 0.1 and each node is allowed up to 10 children. We found that the clustering results on the minigenomes dataset were not altered when these parameters were changed.

### Clustering

An all-against-all pairwise comparison is done on the representative sequences obtained from each Boundary-Tree to construct a distance matrix *S*. For each of the following methods, excluding MCL, and each of the 10 replicate trees, a custom range number of clusters *K* is considered. In the clustering of *S. pneumoniae* pan-genomes, a range of *K*=3000, 3200, 3400, …, 6000 clusters is used.

#### Hierarchical Clustering

an agglomerative hierarchical cluster tree is generated using Ward’s linkage (42) on *S*. Then, the representative sequences are split *into K* clusters.

#### *K*-means Clustering

*S* is clustered using Lloyd’s algorithm (43), with *K*-means++ for cluster center initialization (44). This is an approach to partition sequences into *K* clusters, by iteratively selecting *K* cluster centroids, assigning points to their closest centroids, and updating the centroids based on the new cluster assignments.

*K*-means Vectorized: Since *K*-means is commonly applied to vector data in Euclidean space, we extract from *S*, vectors in Euclidean space. For this, we first generate the symmetric matrix *M*, where 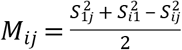. Then, the eigenvalue decomposition *M* = *U V U^T^* is computed, where *U* is orthogonal and *V* is diagonal. The product 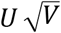 gives Euclidean coordinates for all data points. For the vectorized *K*-means algorithm, we use the same kmeans function, but with 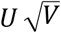 as input instead of S.

#### Spectral Clustering

The distance matrix *S* is transformed into an unweighted adjacency matrix *W* by applying a Gaussian kernel, and thresholding. Then, the graph Laplacian (*L*) and *L*’s eigenvalue decomposition is computed. The top eigenvectors are then clustered using the standard kmeans function. We consider three variants of spectral clustering. One just as described before, which we call SpectralNN, one where *L* is normalized as in in (22), which we call SpectralSM (for Shi-Malik), and one where *L* is normalized as in (23), which we call SpectralNJW (for Ng-Jordan-Weiss).

#### Markov Clustering (MCL)

Similar to Spectral clustering, MCL also uses the adjacency matrix *W*. *W* is column-normalized to yield a stochastic matrix. Then a series of expansion (taking matrix power)-inflation (taking element-wise power)-renormalization steps are applied iteratively on this matrix until the resulting matrix does not change. The nonzero elements of the diagonal correspond to *attractor nodes*. Each attractor, together with all its neighbors in *W* form a cluster (45).

### Error and selection of best number of clusters

In cases where the ground truth is not known, we use the sum of squared errors (SSE) as a measure of clustering quality. SSE is defined as follows:

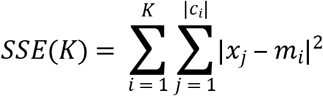

Where *K* is the total number of clusters, *c_i_* is the *i*’th cluster, and |*c_i_*| is the number of elements in *C_i_* . *m_i_* is the medioid (sequence that has the smallest total distance to all other points within the cluster), and *X_j_* is the *j*’th element in *c_i_*.

We compute SSE for a user-defined range of *K* values. The most appropriate number of clusters is determined to be the elbow point, or the point of maximal curvature, of the SSE vs *K* curve. We detect this point by finding the value of *K* where the second derivative of SSE(*K*) is maximized (**Supplementary Figure 2**).

### Consensus clustering

In order to aggregate the replicate Boundary-Forest clustering results, consensus clustering is used (46). First the cluster assignments are extended such that each point that was excluded from the Boundary-Tree gets the cluster assignment of its representative on the tree. This is done for the 10 Boundary-Trees, generating a feature vector of 10 clustering assignments for each sequence, for each clustering method. We then use *K*-medioids clustering, a scalable method, to cluster these feature vectors. For the number of clusters, we use the mode of the best number of clusters from each tree. The feature vectors associated with each sequence is stored for later use, in cluster augmentation.

### Cluster augmentation

Given an existing set of clustered sequences, and a new set of sequences, *cluster augmentation* assigns the new sequences to the closest existing cluster. The new sequences can be processed directly, or the used can choose to do a round of representative selection to reduce the size of the new dataset. A set of representatives is selected from the input sequences by constructing a Boundary-Tree. The representative sequences are then run through the existing Boundary-Forest that was generated when the first set of sequences were clustered. Each representative sequence in the new set traverses each tree in the Boundary-Forest, starting from the root node, by moving to the closest child node. In each tree, the representative is assigned the same cluster as the node it has the smallest Smith-Waterman distance to. This results in as many cluster assignments as the number of trees in the forest. These cluster assignments are taken as a vector, having the same length as the existing feature vector of clustering assignments prior to consensus clustering. The closest existing cluster for each new sequence is determined by 1) finding the vector *v* in the list of stored feature vectors that is closest to the new cluster assignment vector, and 2) assigning to the new sequence the same cluster as that of vector *v*.

### Matching of two clustering partitions

In order to compare two clustering results, or to compare the misclustering error against a known ground truth, we apply the Hungarian matching algorithm (47). Briefly, for clustering *A* and clustering *B*, if we have *n* and *m* clusters respectively, we generate an empty cost matrix *M*: a (*n* + *m*) × (*n* + *m*) matrix of zeros, with each row representing a cluster in *A*, and each column representing a cluster in *B*. The (*i, j*)^th^ entry in this matrix is the dissimilarity cost between cluster *i* from clustering *A* and cluster *j* from clustering *B*. The entries on the upper left *n* × *m* section of *M*, i.e. *M*(1 : *n*, 1 : *m*), are populated with the total number of mismatches between clusters *i* and *j* from clustering *A* and *B* respectively. That is, the sum |*A_i_*\*B_j_*| + |*B_j_*\*A_i_*|, where |*S*| denotes the size of a set *S*. The block *M*(*n*+1 : *n*+*m*, 1 : *m*) represents the costs of clusters in *B* having no representatives in *A*. Each column in this block is populated with |*B_j_*| for cluster *j*. Similarly, *M*(1 : *n*, 1+*m* : *n*+*m*) is populated with |*A_i_*|. Finally, *M*(*n*+1 : *n*+*m*, 1+*m* : *n*+*m*) only has 0 cost. We use this sum of costs to be the error between two clusterings (or a clustering and the ground truth, when the ground truth is known).

### Overlap of two clustering partitions

We define the overlap between clustering partitions *C1* and *C2* on the same dataset as the fraction of clusters in *C1* that are conserved in *C2*. In other words, if a cluster in *C1* has all its members in the same cluster in *C2* (with possibly other sequences included in this cluster in *C2*), it counts towards the overlap. Note that this overlap measure is not symmetrical (i.e. *Overlap*(*C1, C2*) is not necessarily the same as *Overlap(C2, C1)*).

### Confidence scores

We use definitions of item and cluster confidence scores similar to those defined by Monti et al. (26). For a dataset of size *N*, that has been clustered on *T* Boundary-Trees, we define a consensus matrix *M* which is a *N* × *N* matrix, where *M(i,j)* is the proportion of times that items *i* and *j* have appeared in the same cluster. The item consensus for item *i* belonging to cluster *k* is defined as c_i_ 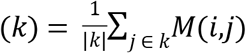 i.e. the average consensus between *i* and other items belonging to the same cluster. Similarly, the cluster consensus for cluster *k* is defined as 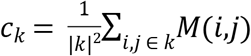 i.e. the average consensus between all pairs of items in custer *k*.

## Acknowledgements

The authors would like to thank Federico Rosconi, Samantha Dyckman and Matt Crum for proofreading, helpful discussions and feedback.

## Supporting Information Legends

**S1 Table.** List of strains used in all 4 *S. pneumoniae* datasets.

**S1 Appendix.** Contains glossary of terms, pseudocode for Boundary-Tree construction, discussion of threshold selection, and Supplementary Figures 1-5.

## References

1. Donati C, Hiller NL, Tettelin H, Muzzi A, Croucher NJ, Angiuoli SV, et al. Structure and dynamics of the pan-genome of Streptococcus pneumoniae and closely related species. Genome Biology. 2010;11:R107.

2. Croucher NJ, Finkelstein JA, Pelton SI, Mitchell PK, Lee GM, Parkhill J, et al. Population genomics of post-vaccine changes in pneumococcal epidemiology. Nat Genet. 2013 Jun;45(6):656–63.

3. van Tonder AJ, Bray JE, Jolley KA, Jansen van Rensburg M, Quirk SJ, Haraldsson G, et al. Genomic Analyses of >3,100 Nasopharyngeal Pneumococci Revealed Significant Differences Between Pneumococci Recovered in Four Different Geographical Regions. Front Microbiol [Internet]. 2019 [cited 2019 May 13];10. Available from: https://www.frontiersin.org/articles/10.3389/fmicb.2019.00317/full

4. Fouts DE, Brinkac L, Beck E, Inman J, Sutton G. PanOCT: automated clustering of orthologs using conserved gene neighborhood for pan-genomic analysis of bacterial strains and closely related species. Nucleic Acids Res. 2012 Dec 1;40(22):e172–e172.

5. Zhao Y, Wu J, Yang J, Sun S, Xiao J, Yu J. PGAP: pan-genomes analysis pipeline. Bioinformatics. 2012 Feb 1;28(3):416–8.

6. Li W, Godzik A. Cd-hit: a fast program for clustering and comparing large sets of protein or nucleotide sequences. Bioinformatics. 2006 Jul 1;22(13):1658–9.

7. Edgar RC. Search and clustering orders of magnitude faster than BLAST. Bioinformatics. 2010 Oct 1;26(19):2460–1.

8. Kavvas ES, Catoiu E, Mih N, Yurkovich JT, Seif Y, Dillon N, et al. Machine learning and structural analysis of Mycobacterium tuberculosis pan-genome identifies genetic signatures of antibiotic resistance. Nature Communications [Internet]. 2018 Dec [cited 2019 Feb 22];9(1). Available from: http://www.nature.com/articles/s41467-018-06634-y

9. Seif Y, Kavvas E, Lachance J-C, Yurkovich JT, Nuccio S-P, Fang X, et al. Genome-scale metabolic reconstructions of multiple Salmonella strains reveal serovar-specific metabolic traits. Nat Commun [Internet]. 2018 Sep 14 [cited 2019 Feb 28];9. Available from: https://www.ncbi.nlm.nih.gov/pmc/articles/PMC6138749/

10. Chaudhari NM, Gupta VK, Dutta C. BPGA-an ultra-fast pan-genome analysis pipeline. Sci Rep [Internet]. 2016 Apr 13 [cited 2019 Feb 28];6. Available from: https://www.ncbi.nlm.nih.gov/pmc/articles/PMC4829868/

11. Ghattargi VC, Gaikwad MA, Meti BS, Nimonkar YS, Dixit K, Prakash O, et al. Comparative genome analysis reveals key genetic factors associated with probiotic property in Enterococcus faecium strains. BMC Genomics [Internet]. 2018 Sep 4 [cited 2019 May 20];19. Available from: https://www.ncbi.nlm.nih.gov/pmc/articles/PMC6122445/

12. Stevens MJA, Tasara T, Klumpp J, Stephan R, Ehling-Schulz M, Johler S. Whole-genomebased phylogeny of Bacillus cytotoxicus reveals different clades within the species and provides clues on ecology and evolution. Sci Rep [Internet]. 2019 Feb 13 [cited 2019 May 20];9. Available from: https://www.ncbi.nlm.nih.gov/pmc/articles/PMC6374410/

13. Zou Y, Xue W, Luo G, Deng Z, Qin P, Guo R, et al. 1,520 reference genomes from cultivated human gut bacteria enable functional microbiome analyses. Nature Biotechnology. 2019 Feb;37(2):179–85.

14. Liu G, Kong Y, Fan Y, Geng C, Peng D, Sun M. Whole-genome sequencing of Bacillus velezensis LS69, a strain with a broad inhibitory spectrum against pathogenic bacteria. Journal of Biotechnology. 2017 May 10;249:20–4.

15. Bhardwaj T, Somvanshi P. Pan-genome analysis of Clostridium botulinum reveals unique targets for drug development. Gene. 2017 Aug 5;623:48–62.

16. Hastie T, Tibshirani R, Friedman J. The Elements of Statistical Learning: Data Mining, Inference, and Prediction, Second Edition. Springer Science & Business Media; 2009. 757 p.

17. Enright AJ, Van Dongen S, Ouzounis CA. An efficient algorithm for large-scale detection of protein families. Nucleic Acids Res. 2002 Apr 1;30(7):1575–84.

18. Ding W, Baumdicker F, Neher RA. panX: pan-genome analysis and exploration. Nucleic Acids Res. 2018 Jan 9;46(1):e5.

19. Page AJ, Cummins CA, Hunt M, Wong VK, Reuter S, Holden MTG, et al. Roary: rapid large-scale prokaryote pan genome analysis. Bioinformatics. 2015 Nov 15;31(22):3691–3.

20. Smith TF, Waterman MS. Identification of common molecular subsequences. Journal of Molecular Biology. 1981 Mar;147(1):195–7.

21. Mathy C, Derbinsky N, Bento J, Rosenthal J, Yedidia J. The Boundary Forest Algorithm for Online Supervised and Unsupervised Learning. arXiv:150502867 [cs, stat] [Internet]. 2015 May 11 [cited 2018 May 25]; Available from: http://arxiv.org/abs/1505.02867

22. Shi J, Malik J. Normalized cuts and image segmentation. IEEE Transactions on Pattern Analysis and Machine Intelligence. 2000 Aug;22(8):888–905.

23. Ng AY, Jordan MI, Weiss Y. On Spectral Clustering: Analysis and an algorithm. In: Dietterich TG, Becker S, Ghahramani Z, editors. Advances in Neural Information Processing Systems 14 [Internet]. MIT Press; 2002 [cited 2019 Jul 5]. p. 849–856. Available from: http://papers.nips.cc/paper/2092-on-spectral-clustering-analysis-and-an-algorithm.pdf

24. Dongen SM van. Graph clustering by flow simulation [Internet]. 2000 [cited 2018 Sep 5]. Available from: http://dspace.library.uu.nl/handle/1874/848

25. Tan P-N, Steinbach M, Karpatne A, Kumar V. Introduction to Data Mining (2Nd Edition). 2nd ed. Pearson; 2018.

26. Monti S, Tamayo P, Mesirov J, Golub T. Consensus Clustering: A Resampling-Based Method for Class Discovery and Visualization of Gene Expression Microarray Data. Machine Learning. 2003 Jul 1;52(1):91–118.

27. Freschi L, Vincent AT, Jeukens J, Emond-Rheault J-G, Kukavica-Ibrulj I, Dupont M-J, et al. The Pseudomonas aeruginosa Pan-Genome Provides New Insights on Its Population Structure, Horizontal Gene Transfer, and Pathogenicity. Genome Biol Evol. 2019 Jan 1;11(1):109–20.

28. Scholz M, Ward DV, Pasolli E, Tolio T, Zolfo M, Asnicar F, et al. Strain-level microbial epidemiology and population genomics from shotgun metagenomics. Nature Methods. 2016 May;13(5):435–8.

29. Raven KE, Reuter S, Gouliouris T, Reynolds R, Russell JE, Brown NM, et al. Genomebased characterization of hospital-adapted Enterococcus faecalis lineages. Nat Microbiol [Internet]. 2016 Mar [cited 2019 Jul 5];1(3). Available from: https://www.ncbi.nlm.nih.gov/pmc/articles/PMC4872833/

30. Schmid M, Muri J, Melidis D, Varadarajan AR, Somerville V, Wicki A, et al. Comparative Genomics of Completely Sequenced Lactobacillus helveticus Genomes Provides Insights into Strain-Specific Genes and Resolves Metagenomics Data Down to the Strain Level. Front Microbiol [Internet]. 2018 [cited 2019 Feb 28];9. Available from: https://www.frontiersin.org/articles/10.3389/fmicb.2018.00063/full

31. Jain M, Olsen HE, Paten B, Akeson M. The Oxford Nanopore MinION: delivery of nanopore sequencing to the genomics community. Genome Biology. 2016 Nov 25;17(1):239.

32. O’Leary NA, Wright MW, Brister JR, Ciufo S, Haddad D, McVeigh R, et al. Reference sequence (RefSeq) database at NCBI: current status, taxonomic expansion, and functional annotation. Nucleic Acids Res. 2016 Jan 4;44(D1):D733–745.

33. Chewapreecha C, Harris SR, Croucher NJ, Turner C, Marttinen P, Cheng L, et al. Dense genomic sampling identifies highways of pneumococcal recombination. Nat Genet. 2014 Mar;46(3):305–9.

34. Cremers AJH, Mobegi FM, de Jonge MI, van Hijum SAFT, Meis JF, Hermans PWM, et al. The post-vaccine microevolution of invasive Streptococcus pneumoniae. Scientific Reports. 2015 Oct 23;5:14952.

35. Jordan IK, Rogozin IB, Wolf YI, Koonin EV. Essential Genes Are More Evolutionarily Conserved Than Are Nonessential Genes in Bacteria.:8.

36. Zhang T, Ramakrishnan R, Livny M. BIRCH: An Efficient Data Clustering Method for Very Large Databases. SIGMOD Rec. 1996 Jun;25(2):103–114.

37. S. Guha, A. Meyerson, N. Mishra, R. Motwani, L. O’Callaghan. Clustering data streams: Theory and practice. IEEE Transactions on Knowledge and Data Engineering. 2003 Jun;15(3):515–28.

38. Cheng S, Karkar S, Bapteste E, Yee N, Falkowski P, Bhattacharya D. Sequence similarity network reveals the imprints of major diversification events in the evolution of microbial life. Front Ecol Evol [Internet]. 2014 [cited 2019 Jul 5];2. Available from: https://www.frontiersin.org/articles/10.3389/fevo.2014.00072/full

39. Sandgren A, Albiger B, Orihuela CJ, Tuomanen E, Normark S, Henriques-Normark B. Virulence in Mice of Pneumococcal Clonal Types with Known Invasive Disease Potential in Humans. J Infect Dis. 2005 Sep 1;192(5):791–800.

40. Wattam AR, Davis JJ, Assaf R, Boisvert S, Brettin T, Bun C, et al. Improvements to PATRIC, the all-bacterial Bioinformatics Database and Analysis Resource Center. Nucleic Acids Res. 2017 Jan 4;45(Database issue):D535–42.

41. Cock PJA, Antao T, Chang JT, Chapman BA, Cox CJ, Dalke A, et al. Biopython: freely available Python tools for computational molecular biology and bioinformatics. Bioinformatics. 2009 Jun 1;25(11):1422–3.

42. Jr JHW. Hierarchical Grouping to Optimize an Objective Function. Journal of the American Statistical Association. 1963 Mar 1;58(301):236–44.

43. Lloyd S. Least squares quantization in PCM. IEEE Transactions on Information Theory. 1982 Mar;28(2):129–37.

44. Arthur D, Vassilvitskii S. K-means++: The Advantages of Careful Seeding. In: Proceedings of the Eighteenth Annual ACM-SIAM Symposium on Discrete Algorithms [Internet]. Philadelphia, PA, USA: Society for Industrial and Applied Mathematics; 2007 [cited 2019 Jul 4]. p. 1027–1035. (SODA ‘07). Available from: http://dl.acm.org/citation.cfm?id=1283383.1283494

45. Van Dongen S. Graph Clustering Via a Discrete Uncoupling Process. SIAM J Matrix Anal Appl. 2008 Jan 1;30(1):121–41.

46. Strehl A, Ghosh J. Cluster Ensembles – A Knowledge Reuse Framework for Combining Multiple Partitions.:35.

47. Munkres J. Algorithms for the Assignment and Transportation Problems. Journal of the Society for Industrial and Applied Mathematics. 1957;5(1):32–8.

